# Larvicidal activity of the photosensitive insecticides, methylene blue and rose bengal, in *Aedes aegypti* and *Anopheles gambiae* mosquitoes

**DOI:** 10.1101/2023.06.30.547250

**Authors:** Cole J. Meier, Julián F. Hillyer

**Affiliations:** Department of Biological Sciences, Vanderbilt University, Nashville, TN, USA

**Author notes:** Corresponding author: Julián F. Hillyer, Ph.D. Department of Biological Sciences Vanderbilt University, VU Station B 35-1634 Nashville, TN 37235.

**Keywords:** Larvicide, Mosquito, Photodynamic, Vector Control, Reactive Oxygen Species, Pesticide Treadmill

## Abstract

**BACKGROUND:** Insecticides are critical for controlling mosquito populations and mitigating the spread of vector–borne disease, but their overuse has selected for resistant populations. A promising alternative to classical chemical insecticides is photosensitive molecules—here called photosensitive insecticides or PSIs—that when ingested and activated by light, generate broadly toxic reactive oxygen species. This mechanism of indiscriminate oxidative damage decreases the likelihood that target site modification-based resistance evolves. Here, we tested whether the PSIs, methylene blue (MB) and rose bengal (RB), are viable insecticides across the mosquito lineage.

**RESULTS:** MB and RB are phototoxic to both *Ae. aegypti* and *An. gambiae* at micromolar concentrations, with greatest toxicity when larvae are incubated in the dark with the PSIs for 2 hr prior to photoactivation. MB is ten times more toxic than RB, and microscopy-based imaging suggests that this is because ingested MB escapes the larval gut and disperses throughout the hemocoel whereas RB remains confined to the gut. Adding food to the PSI-containing water has a bidirectional, concentration-dependent effect on PSI toxicity; toxicity increases at high concentrations but decreases at low concentrations. Finally, adding sand to the water increases the phototoxicity of RB to *Aedes aegypti*.

**CONCLUSION:** MB and RB are larvicidal via a light activated mechanism, and therefore, should be further investigated as an option for mosquito control.

## 1. Introduction

Mosquitoes transmit debilitating diseases such as malaria, dengue, yellow fever, Chikungunya, Zika and lymphatic filariasis, which makes the control of mosquito populations essential for global health.^1–4^ By far the most commonly used method for mosquito control is the application of chemical insecticides,^1–3, 5, 6^ but their persistent use has led to the evolution and selection of resistant mosquito populations.^7–10^ As resistance evolves and spreads, the amount of chemical insecticides required to control mosquito populations increases. Consequently, these insecticides are progressively applied in higher concentrations and continue to accumulate in the environment, which further selects for resistant mosquitoes. This exacerbation of the problem— known as the pesticide treadmill—decreases the efficacy of insecticides against target insects and harms non-target organisms.^11–15^ Therefore, new insecticides that escape this cycle of resistance and environmental accumulation are desperately needed.

Photosensitive insecticides (PSIs) are larvicides that offer an additional insecticide class to work into management rotations.^15^ PSIs are photoactive molecules that generate reactive oxygen species (ROS) in response to light.^16–18^ When ingested by a larva, the ROS produced by the light-stimulated PSIs irreversibly damage adjacent biomolecules. If sufficient damage ensues, the larva dies. Because PSIs damage biomolecules ubiquitously and indiscriminately, target site modification—a primary evolutionary strategy for the development of insecticide resistance ^7, 19^—does not protect larvae against PSIs, although other forms of resistance are possible.^15^

The rapid photodegradation of PSIs is an ecological advantage for these broad-spectrum insecticides. Due to their photoactivation mechanism, PSIs are toxic to small, translucent aquatic insects.^20–26^ However, PSIs are non-toxic to translucent zebrafish even after light irradiation.^27^ The PSIs methylene blue and rose bengal have also been used as therapeutic drugs in humans, including in photodynamic therapy,^28, 29^ which is suggestive of a certain level of safety. The primary environmental advantage of PSIs, however, is that they photodegrade into ecologically harmless byproducts, such as CO_2_, NH_4_^+^, NO_3_^−^, and SO_4_^2–^, in only a few hours.^15, 30, 31^ This is unlike some classical insecticides, which persist in the environment for years.^32–36^

Under laboratory conditions, the PSIs curcumin, rose bengal, hematoporphyrin and eosin-methylene blue are toxic to *Aedes aegypti* larvae, and rose bengal is toxic against *Anopheles gambiae* and *Culex quinquefasciatus.*^20–23^ PSIs are also toxic to Chaoboridae, which is a dipteran family closely related to mosquitoes.^24–26^ However, significant differences in the experimental design of these studies make it difficult to determine which PSIs are most efficient, or how PSI toxicity varies between species. For example, in these studies the larvae of different species were incubated with PSIs at different concentrations, for different durations, and under different feeding regimens. Moreover, the light conditions used to activate the PSIs varied in duration, irradiation intensity, and spectrum. This prevents meaningful comparisons between PSIs and makes it challenging to determine whether a specific PSI is effective against evolutionarily distant mosquito species. Taking a more comprehensive and comparative approach would yield valuable insight into the practical use of PSIs to control mosquito populations.

In this study, we investigated the toxicity of two promising PSIs—methylene blue (MB) and rose bengal (RB)—against *Ae. aegypti* and *An. gambiae*. MB and RB were selected because they have different absorbance spectra and are chemically distinct; MB is a thiazine dye with a molecular weight of 320 g/mol whereas RB is a xanthene dye with a molecular weight of 974 g/mol (Fig. 1A-B). We found that MB and RB are ingested by larvae and become phototoxic at micromolar concentrations, with food and sand altering their lethality. The mosquito lineage (Diptera: Culicidae) originated ∼180 million years ago, and soon thereafter, the two major subfamilies—Culicinae and Anophelinae—diverged.^37, 38^ The finding that MB and RB are toxic to both *Ae. aegypti* (Culicinae) and *An. gambiae* (Anophelinae) indicates that these PSIs are viable candidates for the chemical control of mosquito populations.

**Figure 1.**
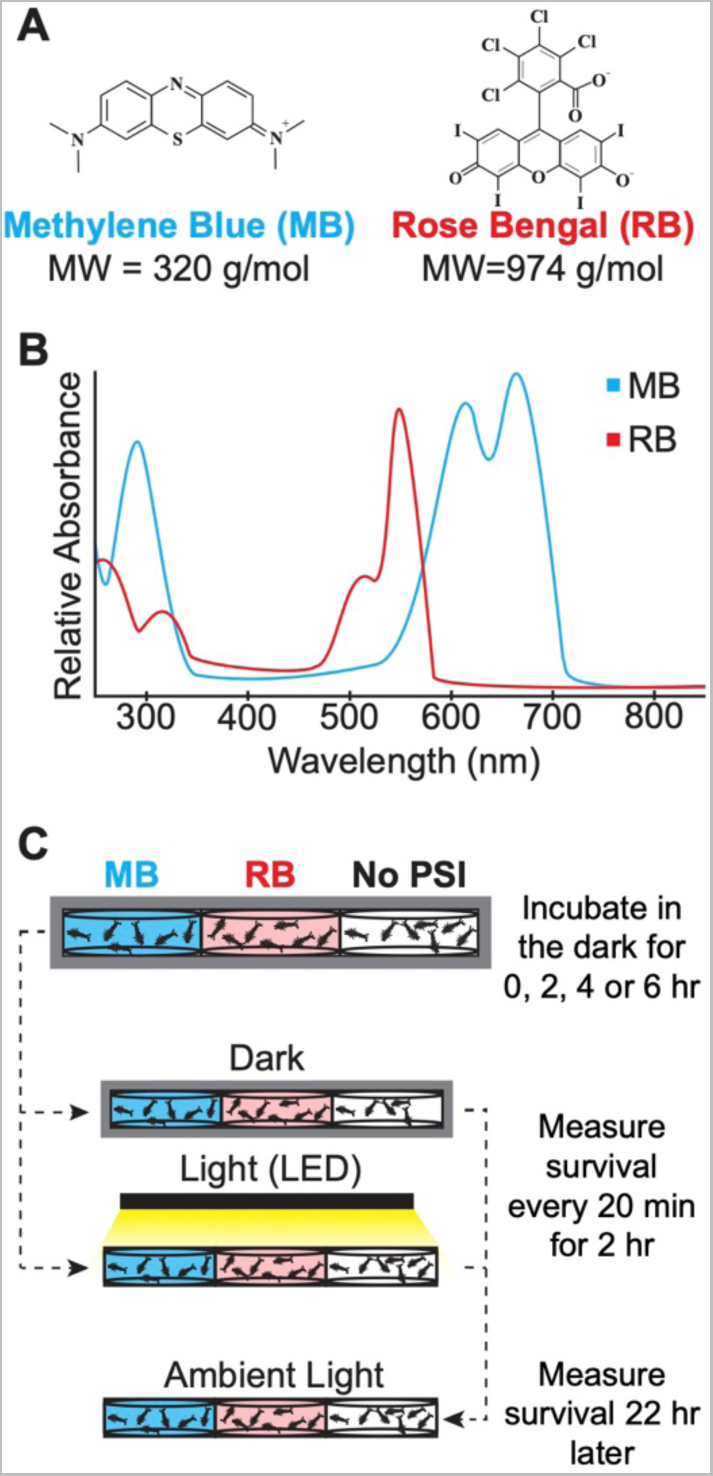
Molecular characteristics of methylene blue and rose bengal, and experimental setup to test larvicidal efficiency. **(A)** Chemical structure and molecular weight of methylene blue and rose bengal. **(B)** Absorbance profile for methylene blue and rose bengal. **(C)** For experiments, larvae were incubated with methylene blue or rose bengal in well plates wrapped in aluminum foil. Larvae were then transferred to a photoperiod (or left in the dark) and survival was measured. MW, molecular weight; MB, methylene blue; RB, rose bengal.

## 2. Materials and Methods

### 2.1 Larval Rearing and Maintenance

*Aedes aegypti* Linnaeus 1762 (Liverpool strain; Diptera: Culicidae) and *Anopheles gambiae* Giles 1902 *sensu stricto* (G3 strain; Diptera: Culicidae) were raised in an environmental chamber under a controlled 12 hr:12 hr ambient light:dark cycle at 27°C and 75% relative humidity. Eggs were hatched in distilled water, and larvae were fed a solution of Wardley Pond Pellets for koi and pond fish (Hartz Mountain Corp., Secaucus NJ) and baker’s yeast (Lesaffre Saf-Instant, Milwaukee WI) daily. All experiments were initiated in 4^th^ instar larvae, and conducted at 27°C.

### 2.2 Incubation of larvae with PSIs, photoactivation, and larval survival

Stock solutions of 2.5 mM of trihydrate methylene blue (Sigma-Aldrich, St. Louis, MO) and disodium salt rose bengal (Thermo Fisher Scientific, Waltham, MA) were prepared by dissolving the chemicals in UV-sterilized, deionized water. Stock solutions were then wrapped in aluminum foil and stored in the dark, at room temperature.

To expose mosquitoes to PSIs, 10 fourth instar larvae were added to each well of a clear 6 well plate. Excess water was removed with a plastic pipette, leaving larvae with only minimal water before immediately adding 5 mL of water containing methylene blue (MB) or rose bengal (RB). The following concentrations of MB and RB were used: MB at 1 µM, 2 µM, 5 µM, 10 µM, and 20 µM; and RB at 2 µM, 5 µM, 10 µM, 20 µM, 50 µM, and 100 µM. These concentrations were determined after a preliminary survey of MB and RB toxicity. As a control, larvae were incubated in water without MB or RB. For all experiments in this study, *Ae. Aegypti* and *An. gambiae* were treated separately; the species were never placed together in a well.

After adding a PSI, a well plate was wrapped in aluminum foil and incubated for 2 hr in the dark at 27°C unless otherwise stated. The lid of the well plate was then removed, and the plate was placed on a white surface that was ∼15 cm directly underneath a 5000 Lumen LED lamp (5000 Lumen LED Work Light, Husky Corporation, Pacific, MO). The time when the lamp was turned on was noted as minute 0 of the photoperiod. Larval survival was then monitored every 20 min for 2 hr, temporarily removing the plate from underneath the lamp for no more than 2 min to count survival. Larval survival was determined via a mechanical stimulus test that used a plastic pipette as a probe; larvae that responded to the stimulus were alive whereas those that did not were dead. Following the completion of the 2 hr photoperiod, the lid was placed back on the well plate and the plate was transferred to the ambient lighting of the environmental chamber. This ambient lighting is insufficiently bright to activate PSIs but maintains the mosquitoes’ diurnal cycle. Larval survival was measured one final time at 22 hr after the photoperiod ended.

All trials were conducted in duplicate well plates, and for each well plate exposed to a photoperiod, an identical well plate was maintained in darkness as a control. These “darkness” plates were sequentially: (i) incubated with PSIs for 2 hr while wrapped in aluminum foil and survival was measured at the end of the incubation (this coincides with darkness incubation of experimental plates), (ii) covered in aluminum foil for another 2 hr and survival was measured again at the end of this second incubation (this coincides with the end of the photoperiod of experimental plates), and (iii) transferred to the ambient light of the environmental chamber and survival was measured one final time 22 hr later (this is similar to what was done for experimental plates that underwent the photoperiod).

### 2.3 Incubation of larvae with PSIs and microscopy of PSI internalization

Larvae were incubated with 5 mL water (no PSI), 20 µM MB, or 100 µM RB at 27°C for 0 hr (∼10 sec), 2 hr, 4 hr, 6 hr, or 8 hr. Larvae were then rinsed in water and transferred to a droplet of water on a microscope slide. Using a Nikon SMZ 1500 stereomicroscope connected to a Nikon Digital Sight DS-Fi1 5 MP CCD Color Camera and Nikon’s Advanced Research NIS Elements software (Nikon Corporation, Tokyo, Japan), larvae were viewed and photographed under oblique coherent contrast trans-illumination.

### 2.4 Incubation of larvae with PSIs plus particulates, photoactivation, and larval survival

Larvae were rinsed 3 times in water to clean them of particulates that were in the colony’s water, and 10 larvae were transferred to each well of a clear 6 well plate. Excess water was removed with a pipette and 5 mL of water was added to every well with a final PSI concentration of 2 µM MB, 20 µM MB, 20 µM RB or 50 µM RB. For each PSI concentration, mosquitoes were separated into three different groups: (i) only PSI in water, (ii) PSI in water plus 100 µL of larval food, or (iii) PSI in water plus 150 mg of sterilized all-purpose sand (American Countryside, Muscle Shoals, AL). Well plates were then wrapped in aluminum foil and incubated for 2 hr at 27°C. Photoactivation and larval survival was then measured as described above, with a matching well plate kept in the dark for every well plate exposed to a photoperiod. Water turbidity in wells with food was similar to water turbidity in wells with sand, as measured spectrophotometrically.

### 2.5 Sampling and statistical analysis

Each treatment was evaluated over 8-10 independent trials, with mosquitoes originating from at least three different egg batches. Data for all trials were combined, and each treatment included 80-100 larvae of each species. Larvae that pupated during the darkness incubation or the photoperiod were excluded from the analysis because pupae do not feed. For each treatment, the data from independent trials were combined, and the median survival (the time at which 50% of larvae had died) and the endpoint survival (the percentage of mosquitoes alive at the end of the experiment) were calculated. Survival curves were compared using the Logrank Mantel Cox test. All data analysis was completed in GraphPad Prism version 9.4.1, and differences were deemed significant at *P* < 0.05.

To evaluate the tissue localization of ingested MB and RB, 30 larvae per species were used for each incubation period and concentration of PSI. Representative images are included in the figures.

## 3. Results

### 3.1 Methylene blue and rose bengal are phototoxic to mosquito larvae, and longer incubations in the dark increase their subsequent phototoxicity

We first sought to determine whether two PSIs—MB and RB—are phototoxic to both *Ae. aegypti* and *An. gambiae*, and how their toxicity is affected by the duration of time the larvae are exposed to the PSI prior to the photoperiod (Fig. 1C). We found that MB and RB are phototoxic to both mosquito species, and that the phototoxicity of both PSIs accelerates when larvae are incubated with the PSI in the dark prior to the initiation of the photoperiod (Fig. 2; Supporting information Table S1). Compared to larvae that were first exposed to the PSI when the photoperiod was initiated, incubating larvae for 2 hr in 20 µM MB decreased the endpoint survival of *Ae. aegypti* from 14% to 0% and of *An. gambiae* from 4% to 0% (Logrank *P* <0.0001 for both species; Fig. 2A-B). Moreover, the median survival of *Ae. aegypti* decreased from 2–24 hr (after the photoperiod, survival was only measured at 2 and 24 hr) to 40 min, and of *An. gambiae* from 100 min to 40 min.

**Figure 2.**
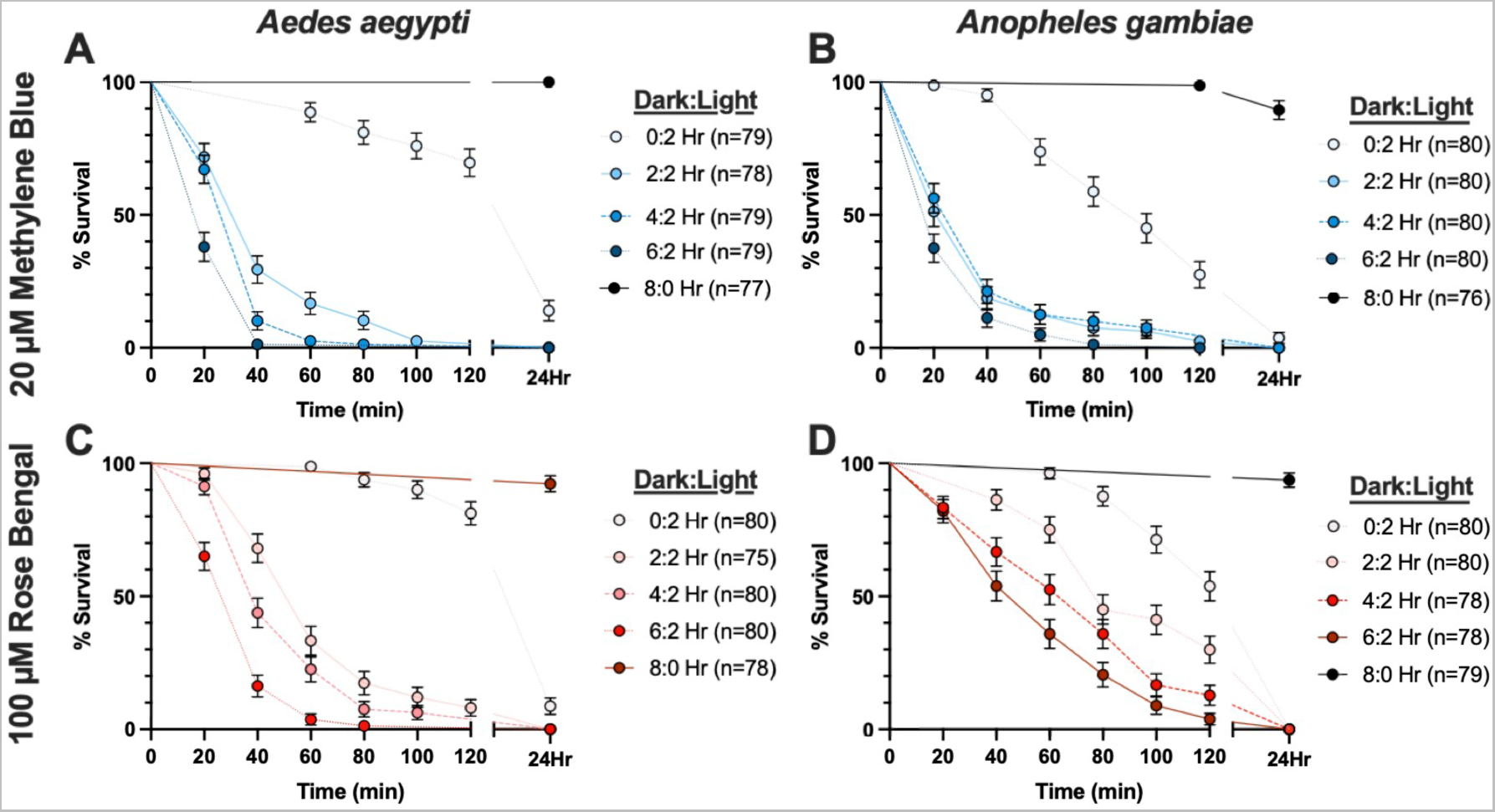
Survival of larvae following various incubation durations with methylene blue and rose bengal. **(A, B)** Survival of *Ae. aegypti* (A) and *An. gambiae* (B) incubated with 20 µM MB in the dark for either 0, 2, 4 or 6 hr and then exposed to either a 2 hr photoperiod or maintained in the dark. **(C, D)** Survival of *Ae. aegypti* (C) and *An. gambiae* (D) incubated with 100 µM RB in the dark for 0, 2, 4 or 6 hr and then exposed to either a 2 hr photoperiod or maintained in the dark. Whiskers indicate the standard deviation, and “n” indicates the number of larvae assayed. Statistical analyses are presented in Supporting information Table S1.

A similar finding was made for RB, where a 2 hr incubation period in 100 µM RB decreased the endpoint survival of *Ae. Aegypti* from 9% to 0%, while the endpoint survival of *An. gambiae* remained at 0% (Logrank *P* <0.0001 for both species; Fig. 2C-D). Moreover, the median survival of *Ae. aegypti* decreased from 2–24 hr to 60 min, and *An. gambiae* from 2–24 hr to 80 min.

When the incubation period in the dark prior to the photoperiod was extended from 2 hr to either 4 hr or 6 hr, MB and RB killed *Ae. aegypti* and *An. gambiae* more quickly. However, because the endpoint survival of all treatments was 0%, we conclude that increasing the incubation period beyond 2 hr does not provide a practical benefit. Finally, to ensure that photoactivation was necessary for PSI toxicity, we incubated larvae in the dark in 20 µM MB or 100 µM RB for 8 hr and then measured survival; in the absence of a photoperiod, all the larvae survived.

Taken altogether, these findings show that MB and RB are both phototoxic to *Ae. aegypti* and *An. gambiae* larvae, and that this phototoxicity accelerates as the darkness incubation duration increases. However, from a practical perspective, phototoxicity is maximized when larvae are incubated with the PSI in the dark for 2 hr prior to light exposure. These findings suggest that PSIs would be most effective when applied between sundown and 2 hr before peak sunlight.

### 3.2 Larvae ingest methylene blue and rose bengal, which localize to different regions of the body

We next used light microscopy to identify whether the accelerated mortality of larvae incubated in MB and RB prior to the onset of the photoperiod was a result of PSI ingestion and accumulation inside the larvae. Based on the coloration of the larvae—with blue indicating MB ingestion and red indicating RB ingestion—ingestion of MB and RB was rapid, but longer incubations increased the ingestion of MB and RB by both *An. gambiae* and *Ae. aegypti* (Fig. 3). For every incubation length, *Ae. aegypti* appeared to ingest slightly more PSI than *An. gambiae*. But more strikingly, MB and RB localized to different regions within the larvae. For both *Ae. aegypti* and *An. gambiae*, MB was found in the gut but also permeated into the hemocoel and dispersed to all regions of the body, whereas RB was confined to the gut (Fig. 3; Fig. 4). This difference in localization is important because photoactivation of a PSI damages molecules in its immediate vicinity. By escaping into the hemocoel, MB should damage the larvae more universally than RB, leading to higher toxicity.

**Figure 3.**
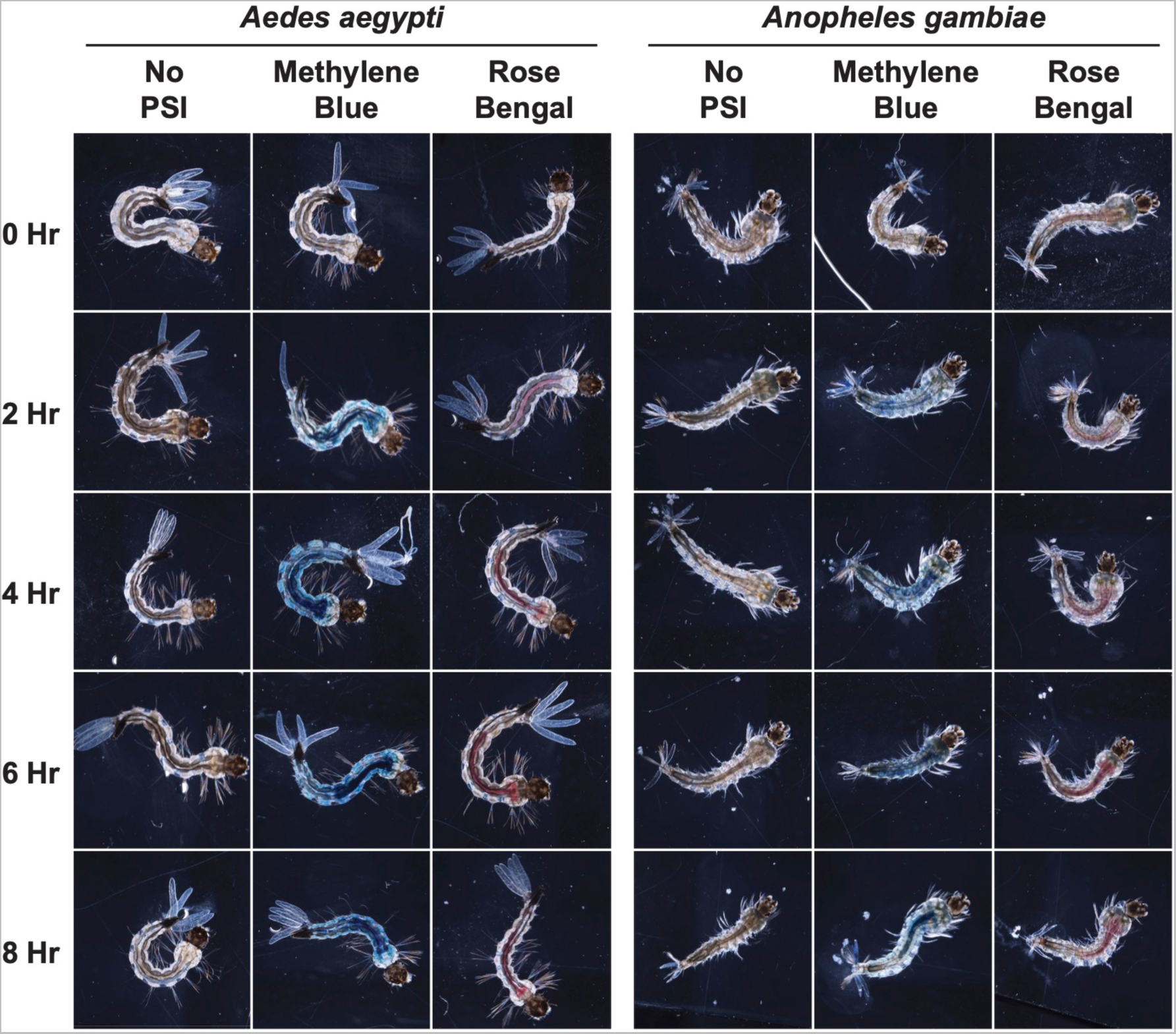
Larval internalization of methylene blue and rose bengal following various incubation durations. Blue indicates methylene blue internalization and red indicates rose bengal internalization.

**Figure 4.**
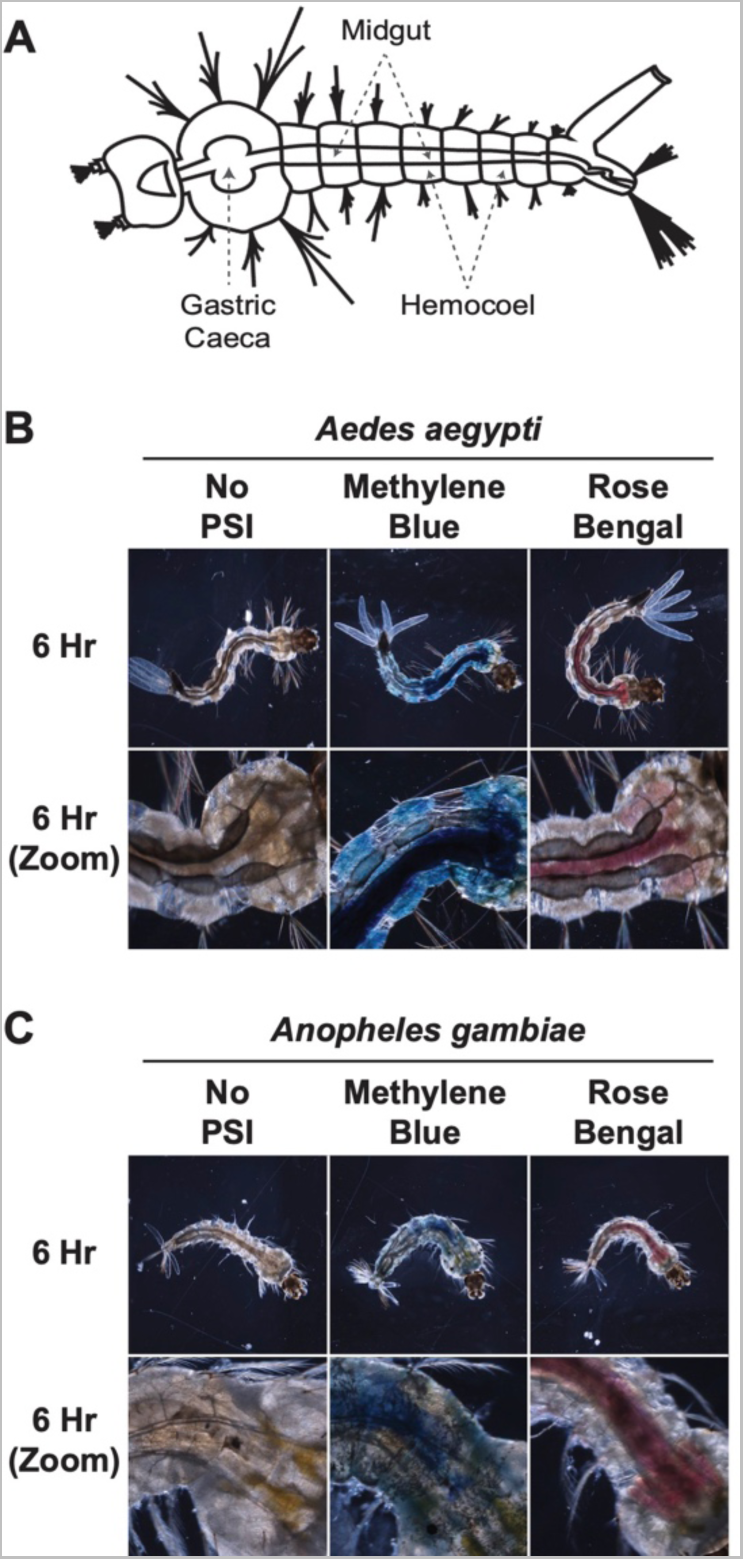
Localization of methylene blue and rose bengal in larvae following a 6 hr incubation. **(A)** Diagram displaying the general anatomy of a mosquito larvae. **(B, C)** *Ae. aegypti* (B) and *An. gambiae* (C) following exposure to methylene blue and rose bengal. Blue indicates methylene blue localization and red indicates rose bengal localization.

### 3.3 Methylene blue and rose bengal are larvicidal at micromolar concentrations

To determine the relative phototoxicity of MB and RB on *Ae. aegypti* and *An. gambiae*, larvae were incubated in the dark for 2 hr with varying concentrations of a PSI and then exposed them to a 2 hr photoperiod (Fig. 1C). As negative controls, larvae were incubated in darkness with these concentrations of PSI but never exposed to a photoperiod.

After exposure to 1 µM MB and a photoperiod, only 33% of *Ae. aegypti* and *An. gambiae* survived by 24 hr (Fig. 5 A-B; Supporting information Table S2). The endpoint and median survival of *Ae. aegypti* and *An. gambiae* continued to decrease with increasing concentrations of PSI until 10 µM MB, where 0% of *Ae. aegypti* and only 2% of *An. gambiae* larvae survived at 24 hr, and the median survival was 40 min (Logrank *P* < 0.0001 for all comparisons). Because 10 µM MB yielded zero or near zero survival, there wasn’t a further increase in mortality when exposing larvae to 20 µM MB (*Ae. aegypti*: *P* = 0.1172; *An. gambiae*: *P* = 0.5847). When larvae were incubated in the dark with MB but not exposed to a photoperiod, the survival of larvae at 24 hr ranged between 93% and 100%. The slight mortality observed was due to larvae predating on one another, and the high survival confirms that photoactivation is required for MB toxicity (Supporting information Fig. S1 A-B; Table S3). Taken altogether, these data show that 10 µM is the lowest concentration of MB that shows the greatest phototoxicity to *Ae. aegypti* and *An. gambiae*, although lower concentrations still provide a high degree of larvicidal activity.

**Figure 5.**
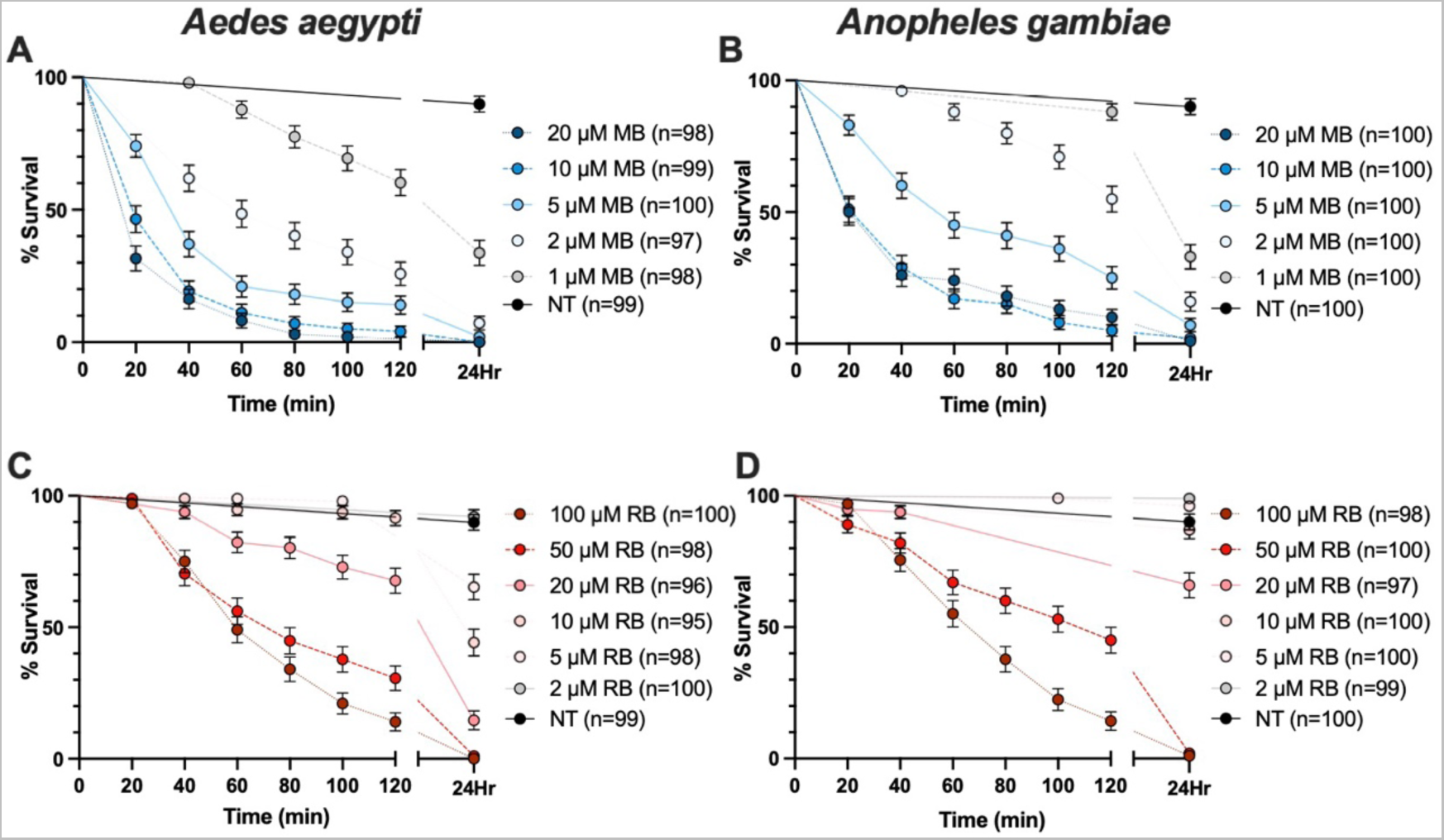
Larval survival following incubation with various concentrations of methylene blue or rose bengal. **(A-D)** Survival of *Ae. aegypti* (A, C) and *An. gambiae* (B, D) following 2 hr incubation and subsequent photoactivation in various concentrations of methylene blue (A, B) and rose bengal (C, D). Whiskers indicate the standard deviation, and “n” indicates the number of larvae assayed. MB, methylene blue; RB, rose bengal; NT, no treatment (no PSI). Statistical analyses are presented in Supporting information Table S2, Table S4, and Table S5.

RB is less toxic than MB. Phototoxicity to *Ae. aegypti* and *An. gambiae* was negligible until the concentration reached 5 µM and 10 µM, respectively (*Ae. aegypti*: *P* < 0.0001; *An. gambiae*: *P* = 0.0248; Fig. 5 C-D; Supporting information Table S2). At 5 µM RB, 65% and 96% of *Ae. aegypti* and *An. gambiae* survived by 24 hr, respectively. At 10 µM RB, survival decreased further, with only 44% of *Ae. aegypti* and 87% of *An. gambiae* surviving by 24 hr. At higher concentrations of RB, endpoint and median survival continued to decrease until 50 µM, where only 1% of *Ae. aegypti* and 2% of *An. gambiae* survived by 24 hr, and the median survival was 80 min and 120 min, respectively (*P* < 0.0001 for both species). At 100 µM RB, the median survival decreased further, but because the endpoint survival only decreased by 1% for *Ae. aegypti* and *An. gambiae*, we conclude that increasing RB concentration beyond 50 µM does not meaningfully increase toxicity. When larvae were incubated in the dark with RB but not exposed to a photoperiod, survival was between 92% and 100%. The slight mortality observed was due to larvae predating on one another, and the high survival confirms that photoactivation is required for RB toxicity (Supporting information Fig. S1 C-D; Table S3). Taken altogether, these data show that 50 µM is the lowest concentration of RB that shows the greatest toxicity to *Ae. aegypti* and *An. gambiae*, and that concentrations below 50 µM RB are largely ineffective.

When we compared the effect of equivalent concentrations of MB and RB on larval survival, we found that MB was more toxic than RB (Fig. 5; Supporting information Table S4). For example, when *Ae. aegypti* were exposed to 2 µM of each PSI, 7% survived by 24 hr when exposed to MB whereas 92% survived when exposed to RB (*P* < 0.0001). Likewise, when *An. gambiae* were exposed to 2 µM of each PSI, 16% survived by 24 hr when exposed to MB whereas 99% survived when exposed to RB (*P* < 0.0001).

When we compared the likelihood that *Ae. aegypti* and *An. gambiae* would survive a PSI exposure, we found that *Ae. aegypti* are slightly more susceptible to PSI toxicity than *An. gambiae* (Fig. 5; Supporting information Table S5). At 2 µM MB, for example, 7% of *Ae. aegypti* and 16% of *An. gambiae* survived by 24 hr (*P* < 0.0001), and the median survival for these two species was 60 min and 2–24 hr, respectively. Overall, these data show that MB is a more efficient larvicide than RB, and that while both PSIs are effective, MB and RB are more phototoxic against *Ae. aegypti*.

### 3.4 Larval food has a bidirectional, concentration-dependent effect on phototoxicity

Based on the outcomes of concentration-dependent survival curves and microscopy-based examinations of larvae, we (i) inferred that PSI phototoxicity requires PSI ingestion and (ii) speculated that stimulating feeding enhances PSI toxicity. Because the phototoxicity of curcumin to *Culex pipiens* is enhanced when it is mixed with D-mannitol but the toxicity of RB against *Culex quinquefasciatus* decreases when applied to cesspits,^20, 39^ we tested whether stimulating feeding by exposing larvae to a mixture of PSI and food enhances toxicity. For this, larvae were incubated for 2 hr in the dark with a PSI plus (i) larval food, (ii) sand, or (iii) no added particulates (solution). Sand was used to create an environment with equal turbidity to that of larval food, but using particulates that are non-nutritious and less likely to stimulate feeding. For the PSIs, the concentrations were chosen because they exhibit intermediate (2 µM MB and 20 µM RB) and high (20 µM MB and 50 µM RB) toxicity during a photoperiod. After the incubation, larvae were exposed to a 2 hr photoperiod and survival monitored every 20 min throughout the photoperiod, and again 22 hr later. Without any PSI, neither sand nor food affected larval survival in either light or dark conditions, indicating that neither food nor sand are toxic (Supporting information Fig. S2; Table S6). Similarly, in the presence of either PSI but in the absence of a photoperiod, larval survival was unaffected by the addition of food or sand, indicating that any toxicity would be due to PSI photoactivation (Supporting information Fig. S3; Table S7; Fig. S4; Table S8).

We began these experiments by testing MB (Fig. 6; Supporting information Table S9). As expected given our earlier findings, exposure to 2 µM and 20 µM MB in the absence of a particulate (food or sand) resulted in intermediate and high toxicity following a photoperiod. Adding food to 2 µM MB decreased phototoxicity; food increased the endpoint survival of *Ae. aegypti* from 4% to 21%, and of *An. gambiae* from 19% to 59% (Logrank *P* < 0.0001 for both species). Likewise, adding food to 2 µM MB increased the median survival of *Ae. aegypti* from 60 min to 110 min, and of *An. gambiae* from 2–24 hr to >24 hr (>50% survived when food was added). We observed the opposite effect at 20 µM MB, where food increased phototoxicity; food decreased the endpoint survival of *An. gambiae* from 15% to 1%, but for *Ae. aegypti* we could not detect a change because only 1% of larvae survived when exposed to MB alone or with the addition of food. However, adding food to 20 µM MB decreased the median survival of *Ae. aegypti* from 40 min to 20 min and of *An. gambiae* from 120 min to 40 min (Logrank *P* < 0.05 for both species). Adding sand to the water did not affect MB-mediated phototoxicity, indicating that the changes are due to food and not water turbidity. Taken altogether, adding food has a bidirectional effect on MB toxicity; food decreases phototoxicity at a low MB concentration but increases phototoxicity at a high MB concentration (Fig. 6; Supporting information Table S9).

**Figure 6.**
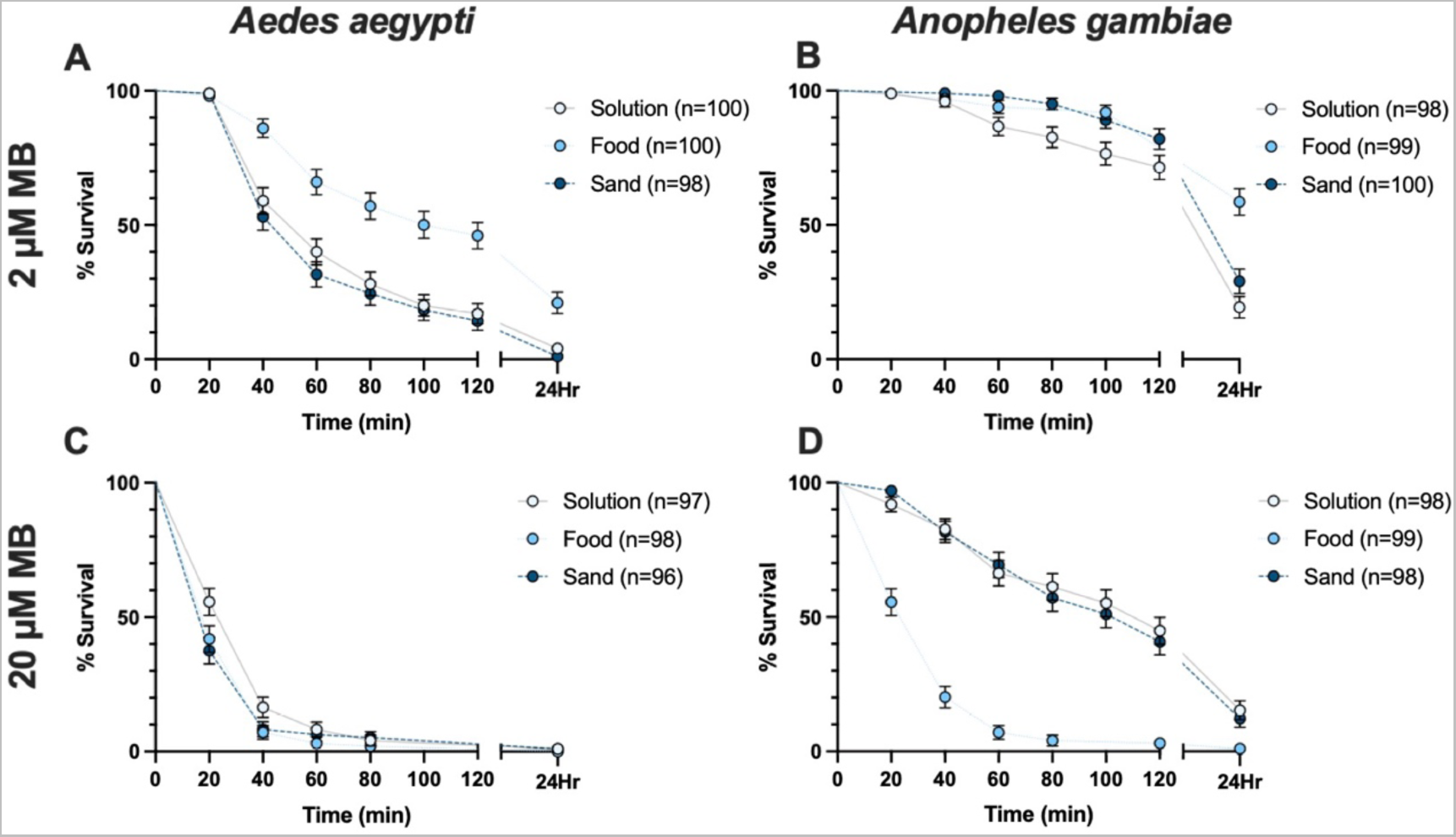
Survival of larvae incubated with methylene blue in the presence of sand or food. **(A-D)** Survival of *Ae. aegypti* (A, C) and *An. gambiae* (B, D) following a 2 hr incubation and subsequent photoactivation in 2 µM methylene blue (A, B) or 20 µM methylene blue (C, D) without particulates (solution), with food, or with sand. Whiskers indicate the standard deviation, and “n” indicates the number of larvae assayed. Statistical analyses are presented in Supporting information Table S9.

We then turned our attention to RB (Fig. 7; Supporting information Table S10). As expected given our earlier findings, exposure to 20 µM and 50 µM RB in the absence of a particulate resulted in intermediate and high toxicity following a photoperiod. Adding food altered RB-mediated phototoxicity, but in a manner different from MB. When larvae were exposed to 20 µM RB, adding food decreased the endpoint survival of *Ae. aegypti* from 35% to 0% (Logrank *P* < 0.0001) but left the endpoint survival of *An. gambiae* unchanged (39% versus 32%; *P* = 0.6149). Likewise, adding food decreased the median survival of *Ae. aegypti* from 2– 24 hr to 40 min, whereas the median survival of *An. gambiae* remained at 2–24 hr—although there was a trend for accelerated mortality in the presence of food. At 50 µM RB alone, only 3% of *Ae. aegypti* and 2% of *An. gambiae* survived at 24 hr. Adding food did not alter endpoint survival likely because the low survival following 50 µM RB alone meant that we could not meaningfully detect whether the addition of food decreased endpoint survival further. However, adding food to 50 µM RB decreased the median survival of *Ae. aegypti* from 80 min to 40 min and of *An. gambiae* from 100 min to 80 min (Logrank *P* < 0.05 for both species).

**Figure 7.**
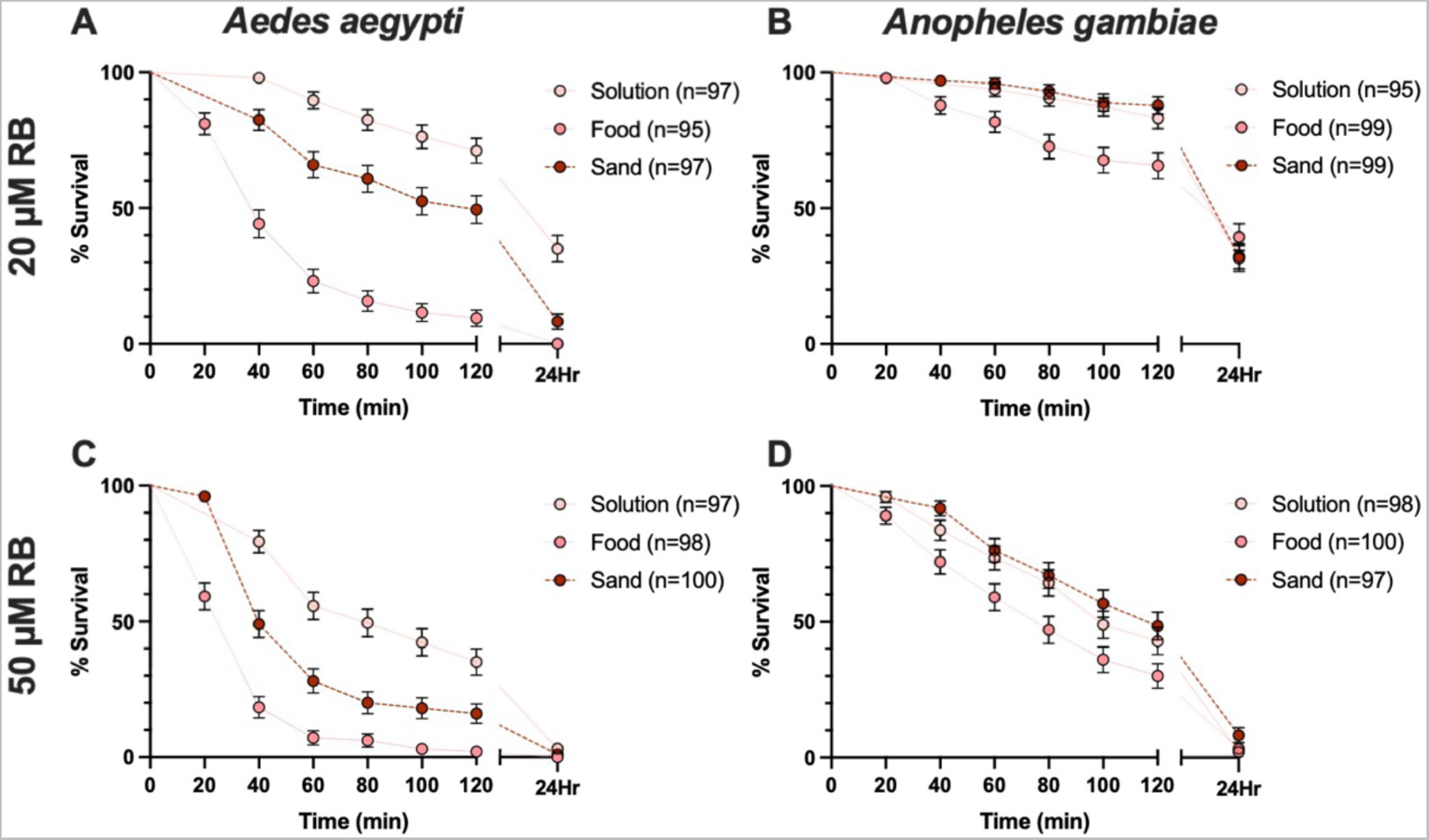
Survival of larvae incubated with rose bengal in the presence of sand or food. **(A-D)** Survival of *Ae. aegypti* (A, C) and *An. gambiae* (B, D) following a 2 hr incubation and subsequent photoactivation in 20 µM rose bengal (A, B) or 50 µM rose bengal (C, D) without particulates (solution), with food, or with sand. Whiskers indicate the standard deviation, and “n” indicates the number of larvae assayed. Statistical analyses are presented in Supporting information Table S10.

Surprisingly, adding sand to RB decreased the endpoint and median survival of *Ae. aegypti* but not *An. gambiae*. At 20 µM RB, adding sand decreased the endpoint survival of *Ae. aegypti* from 35% to 8%, and the median survival from 2–24 hr to 120 min (Logrank *P* < 0.0001). At 50 µM RB alone, only 3% of *Ae. aegypti* survived by 24 hr, so any detrimental effect that sand may have on endpoint survival could not be detected. However, adding sand to 50 µM RB decreased the median survival of *Ae. aegypti* from 80 min to 40 min (Logrank *P* < 0.0001). Taken altogether, adding food increases the phototoxicity of RB against *Ae. aegypti* and *An. gambiae,* with a more pronounced effect at higher PSI concentrations (Fig 7; Supporting information Table S10). Furthermore, adding sand increased the toxicity of RB against *Ae. aegypti* but not *An. gambiae*.

## 4. Discussion

We determined the larvicidal efficiency of two promising PSIs on two evolutionarily distant mosquito species that transmit disease to humans and animals. We found that (i) the ideal time to apply a PSI is at least 2 hr prior to peak sunlight, (ii) MB—which is smaller than RB—is more toxic than RB likely because it disperses more broadly throughout the larvae after being ingested and therefore can damage tissues more diversely, (iii) *Ae. aegypti* are slightly more susceptible to PSIs than *An. gambiae,* (iv) the presence of food in the PSI solution can bidirectionally modulate PSI toxicity in a concentration-dependent manner, and (v) the presence of sand can affect phototoxicity. Overall, these findings suggest that PSIs are viable candidates for the control of mosquito populations (Fig. 8).

**Figure 8.**
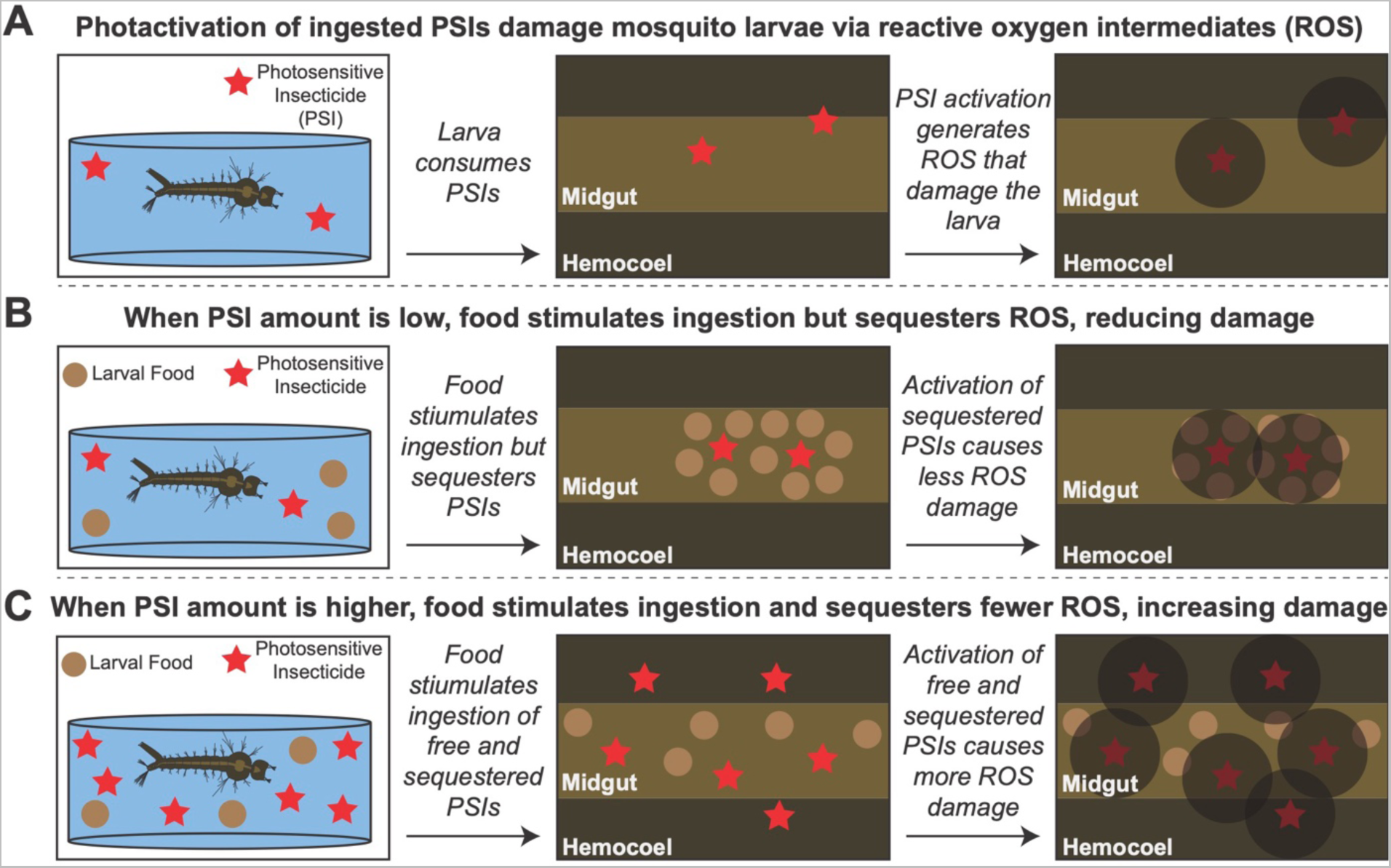
Model of PSI toxicity, and the effects that larval food has on PSI toxicity. **(A)** Model of PSI toxic mechanism in the absence of food. **(B)** When PSI amount is low, food stimulates ingestion of PSIs but sequesters the PSIs and ROS within the larvae, thereby reducing damage. **(C)** When PSI amount is higher, food stimulates the ingestion of PSIs and sequesters fewer ROS, thereby increasing ROS-mediated damage.

Considering how rapidly they photodegrade,^15^ PSIs are presumably more effective when applied before peak sunlight to allow for sufficient ingestion prior to photoactivation. Yet, how long before peak sunlight a PSI should be applied to best control mosquito populations remained unknown. Investigators recently assessed how the absorbance of a PSI solution changes after having been incubated with *Culex pipiens* larvae, and suggested that a 6 hr incubation period in darkness is ideal.^40^ However, this study did not empirically test whether this incubation—relative to any other incubation—results in maximum toxicity. Other investigators measured how the PSI, chlorophyllin, accumulates in the glass worm larvae, *Chaoborus crystallinus*, and found that 2–4 ng of chlorophyllin accumulates per hour.^25^ Under this scenario, a 3 hr incubation was needed to achieve optimal phototoxicity.^26^ Although glassworm larvae share many morphological similarities with mosquito larvae, we predict that the ideal time of PSI application is affected by the feeding behavior of the target species, so we empirically tested this for mosquitoes. We observed that a 2 hr incubation with MB or RB in the dark prior to a photoperiod is sufficient to kill nearly all the larvae of both *Ae. aegypti* and *An. gambiae*. Together with the observations that significant mortality occurs without a preincubation in the dark and that incubations that last longer than 2 hr result in total mortality, these findings demonstrate that the window of PSI application for mosquito control is flexible, but that the ideal time of application is between sundown and 2 hr before peak sunlight the following day.

Once ingested, it is unclear which factors determine how PSIs disperse within a larva and whether their localization affects phototoxicity. When the PSI curcumin is ingested by *Ae. aegypti* larvae, for example, it accumulates throughout the gut and permeates into the hemocoel,^39^ but when the PSIs TPPS, TMPyP and mTHPC (porphyrin derivatives) are ingested by *Chaoborus crystallinus* larvae, they all remain confined to the larval gut.^24^ Here, we observed that MB—which is similar in size to curcumin—accumulates in the gut, gastric caeca and hemocoel of both *Ae. aegypti* and *An. gambiae*. In contrast, RB—which is similar in size to TPPS, TMPyP and mTHPC, and therefore much larger than curcumin or MB—remains confined to the gut and gastric caeca. From this, we infer that smaller PSIs can cross the midgut epithelium and spread throughout the body. It is possible that other molecular characteristics such as hydrophobicity or charge may also play a role in determining the PSI’s localization. However, given that TPPS, TMPyP and mTHPC (i) each hold a different charge—anionic (−4), neutral, and cationic (+4), respectively—and (ii) vary in their hydrophobicity, we conclude that molecular size is the most likely determinant of PSI localization.

In this study we conducted parallel experiments in *Ae. aegypti* and *An. gambiae*, which allowed us to compare the efficacy of two PSIs in evolutionarily distant mosquito species. We found that both MB and RB are viable insecticides against *Ae. aegypti* and *An. gambiae*, with the endpoint mortality (at 24 hr) at a given PSI concentration usually being similar for both species. Although the endpoint mortality was similar, *Ae. aegypti* died more quickly from PSI phototoxicity than *An. gambiae*, as seen in the slopes of the survival curves. We suspect that the higher vulnerability of *Ae. aegypti*, despite this mosquito being larger and slightly darker than *An. gambiae*, is due to increased PSI ingestion.

When curcumin is mixed with sucrose and D-mannitol, its phototoxicity against *Culex pipiens* larvae increases.^39^ This suggests that larval feeding affects PSI toxicity. However, the organic matter that larvae naturally ingest is more complex than pure sugar; larvae ingest a combination of carbohydrates, proteins and lipids, all as part of cellular and environmental debris. The presence of complex foods in the gut at the time of PSI ingestion may affect toxicity in several ways, including by (i) reducing translucency and thereby decreasing toxicity, (ii) sequestering free PSI and thereby decreasing toxicity, or (iii) stimulating feeding thereby increasing toxicity. When applied to cesspits with an abundance of organic matter, RB phototoxicity against *Culex quinquefasciatus* decreases dramatically.^20^ Therefore, we investigated how organic and inorganic particulates in the water affect PSI toxicity. Interestingly, adding larval food (consisting of baker’s yeast and koi food) causes a bidirectional effect on PSI phototoxicity that depends on the concentration of the PSI (Fig. 8). At a high PSI concentration, food increases phototoxicity whereas at a low PSI concentration, and more specifically a low concentration of MB, food decreases phototoxicity. We suspect that PSIs bind to the larval food, resulting in more PSI being ingested. At a low PSI concentration, we predict that the ROS produced within the larvae upon photoactivation react primarily with the food instead of larval tissues, but at a high PSI concentration, we predict that there remains enough PSI in solution such that the ROS cause damage to larval tissues. In the case of MB, this occurs both within the midgut and the hemocoel. This explains why at 2 µM MB (a low concentration) adding food decreases toxicity whereas at 20 µM RB, 20 µM MB or 50 µM RB (high concentrations) adding food increases the toxicity. We further predict that the bidirectional effect on PSI phototoxicity depends on the ratio of organic matter to PSI concentration. If applied properly, the addition of larval food—or the larval food already present in breeding areas—could bolster the larvicidal efficacy of PSIs. However, larvae in nature feed on a much more complex array of food and the environment is more complex, so additional studies in more natural settings, like mesocosms, are needed to understand this interaction more fully.

Interestingly, adding sand, which sequesters PSIs and over several hours settles to the bottom, increased the toxicity of both MB and RB to *Ae. aegypti* yet had a negligible effect on *An. gambiae.* This species-specific effect is likely related to the feeding behavior of these species. Anopheline larvae are primarily surface feeders whereas culicine larvae switch between surface feeding and submerged feeding.^41^ Therefore, *Ae. aegypti* larvae are likely to graze on or consume sand that has bound PSIs, whereas *An. gambiae* larvae are not. Silicon oxide (SiO_2_) is a major component of sand, and in nanoparticle form enhances the delivery of photosensitive molecules for photodynamic therapy.^42^ Therefore, encapsulating PSIs in organic or inorganic matter before application may increase PSI efficacy. Yeast encapsulation of larvicidal essential oils increased their efficacy while also providing benefits to the surrounding microbial environment, which encounters the vehicle rather than the toxic package.^43, 44^ Encapsulating PSIs in yeast may have a similar, positive effect.

Our experiments were conducted in the laboratory using artificial lighting. It is likely that MB and RB are more toxic when irradiated by sunlight instead of our 5000 lumen LED lamp because sunlight has a more uniform emission spectra and is ∼50 times brighter than our lamp. When it comes to their individual effectiveness, MB is more toxic than RB, which suggests that smaller PSIs that disperse throughout the hemocoel are more toxic than larger PSIs that remain confined to the gut. Moreover, MB likely has the practical advantage that by virtue of being smaller it disperses more quickly in a water column if it were used in a larger body of water. We predict that MB and RB would be even more effective in combination because they have different absorbance spectral profiles; by absorbing photons at different wavelengths, these PSIs do not compete with one another for the production of ROS (Fig. 1). Under these optimal conditions, additional studies that explore the relative toxicity of PSIs compared with other larvicides are needed.

## 5. Conclusion

MB and RB are inexpensive chemicals that have a long shelf-life when they are stored in the dark or highly concentrated. Here, we show that they are larvicidal via a light-activated mechanism at micromolar concentrations. Because MB and RB kill larvae via indiscriminate oxidative damage that is less likely to select for resistance, and because they rapidly photodegrade and thereby are expected to have negligible detrimental effects on ecosystems,^15^ MB and RB are promising larvicides that should be further investigated as an option for the control of mosquito populations.

## Abbreviations

PSI: Photosensitive insecticide
ROS: Reactive oxygen species
NT: No treatment
MB: Methylene blue
RB: Rose bengal
*An. gambiae*: *Anopheles gambiae*
*Ae. aegypti*: *>Aedes aegypti.*

## Acknowledgements

We thank Jordyn Barr and Lindsay Martin for helpful feedback on the manuscript. Useful discussions with Dr. Matthew Rouhier are greatly appreciated. This research was funded by Vanderbilt University institutional funds.

## Data Sharing and Data Accessibility

The datasets supporting the conclusions of this article are included within the article and its supplementary files. This includes Supporting information: Data, where all the survival data are presented in the format for analysis in GraphPad Prism.

## Authors’ contributions

CJM and JFH designed the study; CJM performed the experiments; CJM and JFH analyzed the data and wrote the manuscript.

## Supplementary Information

**Table S1.** Descriptive statistics of larval survival following various incubation periods with methylene blue and rose bengal.

**Table S2.** Descriptive statistics of larval survival following incubation and photoperiod of various concentrations of PSIs.

**Table S3.** Descriptive statistics of larvae incubated with various concentrations of PSIs in the dark.

**Table S4.** Comparison of larval survival between methylene blue and rose bengal following a photoperiod.

**Table S5.** Comparison of *Aedes aegypti* and *Anopheles gambiae* survival following incubation with PSIs and a photoperiod.

**Table S6.** Descriptive statistics of larvae following incubation in water and particulates.

**Table S7.** Descriptive statistics of larvae following incubation in methylene blue and particulates, without a photoperiod.

**Table S8.** Descriptive statistics of larvae following incubation in rose bengal and particulates, without a photoperiod.

**Table S9.** Descriptive statistics of larvae following incubation in methylene blue and particulates, with a photoperiod.

**Table S10.** Descriptive statistics of larvae following incubation in rose bengal blue and particulates, with a photoperiod.

**Figure S1.** Larval survival following incubation with various concentrations of PSIs in the dark.

**Figure S2.** Larval survival in the presence of sand or food in the dark.

**Figure S3.** Survival of larvae incubated in methylene blue and particulates, without a photoperiod.

**Figure S4.** Survival of larvae incubated in rose bengal and particulates, without a photoperiod.

**Data.** Survival data, presented in the format for analysis in GraphPad Prism.

