## Supplementary material for "Larvicidal activity of the photosensitive insecticides, methylene blue and rose bengal, in *Aedes aegypti* and *Anopheles gambiae* mosquitoes": Meier and Hillyer 2023 bioRxiv Supporting Info.pdf

<ins>Supplementary tables and figures:</ins>

- Tables S1-S10
- Figures S1-S4

**Table S1. Descriptive statistics of larval survival following various incubation periods with methylene blue and rose bengal.** Larval survival was measured following an 8 hr incubation with either 20 µM MB or 100 µM RB in the dark. Additionally, larval survival was measured following a 0, 2, 4, or 6 hr incubation in the dark with either 20 µM MB or 100 µM RB and then a 2 hr photoperiod. Survival was compared using the Log-Rank Mantel Cox test; all significance indicates an increase in toxicity from the previous incubation time:photoperiod.

| | *Aedes aegypti* | | | | *Anopheles gambiae* | | | |
|---|---|---|---|---|---|---|---|---|
| **20 µM Methylene blue** | | | | | | | | |
| **Incubation Time: Photoperiod** | **Median Survival ‡** | **Endpoint Survival** | **n** | **Logrank *P* ¶** | **Median Survival ‡** | **Endpoint Survival** | **n** | **Logrank *P* ¶** |
| 8 hr: 0 hr | N/A | 100% | 77 | N/A | N/A | 89% | 76 | N/A |
| 0 hr: 2 hr | 24 hr | 14% | 79 | **<0.0001** | 100 min | 4% | 80 | **<0.0001** |
| 2 hr: 2 hr | 40 min | 0% | 78 | **<0.0001** | 40 min | 0% | 80 | **<0.0001** |
| 4 hr: 2 hr | 40 min | 0% | 79 | **0.0103** | 40 min | 0% | 80 | 0.3651 |
| 6 hr: 2 hr | 20 min | 0% | 79 | **<0.0001** | 20 min | 0% | 80 | **0.0034** |
| **100 µM Rose bengal** | | | | | | | | |
| **Incubation Time: Photoperiod** | **Median Survival ‡** | **Endpoint Survival** | **n** | **Logrank *P* ¶** | **Median Survival ‡** | **Endpoint Survival** | **n** | **Logrank *P* ¶** |
| 8 hr: 0 hr | N/A | 92% | 78 | N/A | 60 min | 0% | 79 | **0.0179** |
| 0 hr: 2 hr | 24 hr | 9% | 80 | **<0.0001** | N/A | 94% | 80 | N/A |
| 2 hr: 2 hr | 60 min | 0% | 75 | **<0.0001** | 24 hr | 0% | 80 | **<0.0001** |
| 4 hr: 2 hr | 40 min | 0% | 80 | **0.0281** | 80 min | 0% | 78 | **<0.0001** |
| 6 hr: 2 hr | 40 min | 0% | 80 | **<0.0001** | 80 min | 0% | 78 | **0.0009** |

‡ N/A – not applicable because survival exceeded 50%

¶ P<0.05 are bolded

**Table S2. Descriptive statistics of larval survival following incubation and photoperiod of various concentrations of PSIs.** Larval survival was measured following a 2 hr incubation with 1 µM, 2 µM, 5 µM, 10 µM or 20 µM of methylene blue, or 2 µM, 5 µM, 10 µM, 20 µM, 50 µM or 100 µM of rose bengal, followed by a photoperiod. The Log-Rank Mantel Cox test was used to compare the survival between each prior dose of the same PSI and species. Statistical significance indicates an increase in toxicity.

| | ***Aedes aegypti*** | | | | ***Anopheles gambiae*** | | | |
|---|---|---|---|---|---|---|---|---|
| **Methylene blue** | | | | | | | | |
| **Concentration** | **Median Survival ‡** | **Endpoint Survival** | **n** | **Logrank *P* ¶** | **Median Survival ‡** | **Endpoint Survival** | **n** | **Logrank *P* ¶** |
| NT | N/A | 100% | 99 | N/A | N/A | 90% | 100 | N/A |
| 1 µM | 24 hr | 33% | 98 | **<0.0001** | 24 hr | 33% | 100 | **<0.0001** |
| 2 µM | 60 min | 7% | 97 | **<0.0001** | 24 hr | 16% | 100 | **<0.0001** |
| 5 µM | 40 min | 2% | 100 | **<0.0001** | 60 min | 7% | 100 | **<0.0001** |
| 10 µM | 20 min | 0% | 99 | **<0.0001** | 40 min | 2% | 100 | **<0.0001** |
| 20 µM | 20 min | 0% | 98 | 0.1172 | 30 min | 1% | 100 | 0.5837 |
| **Rose bengal** | | | | | | | | |
| **Concentration** | **Median Survival ‡** | **Endpoint Survival** | **n** | **Logrank *P* ¶** | **Median Survival ‡** | **Endpoint Survival** | **n** | **Logrank *P* ¶** |
| NT | N/A | 90% | 99 | N/A | N/A | 90% | 100 | N/A |
| 2 µM | N/A | 92% | 100 | 0.6063 | N/A | 99% | 99 | 0.0056 |
| 5 µM | N/A | 65% | 98 | **<0.0001** | N/A | 96% | 100 | 0.1778 |
| 10 µM | 24 hr | 44% | 95 | **0.0018** | N/A | 87% | 100 | **0.0248** |
| 20 µM | 24 hr | 15% | 96 | **<0.0001** | N/A | 66% | 97 | **0.0003** |
| 50 µM | 80 min | 1% | 98 | **<0.0001** | 120 min | 2% | 100 | **<0.0001** |
| 100 µM | 60 min | 0% | 100 | **0.0208** | 80 min | 1% | 98 | **<0.0001** |

‡ N/A – not applicable because survival exceeded 50%
¶ P<0.05 are bolded

**Table S3. Descriptive statistics of larvae incubated with various concentrations of PSIs in the dark.** Larval survival was measured following a 2 hr incubation with 1 µM, 2 µM, 5 µM, 10 µM or 20 µM of methylene blue, or 2 µM, 5 µM, 10 µM, 20 µM, 50 µM or 100 µM of rose bengal, without a photoperiod. The Log-Rank Mantel Cox test was used to compare the survival between each prior dose of the same PSI and species. Statistical significance indicates an increase in toxicity.

| | *Aedes aegypti* | | | | *Anopheles gambiae* | | | |
|---|---|---|---|---|---|---|---|---|
| **Methylene blue** | | | | | | | | |
| **Concentration** | **Median Survival ‡** | **Endpoint Survival** | **n** | **Logrank *P* ¶** | **Median Survival ‡** | **Endpoint Survival** | **n** | **Logrank *P* ¶** |
| NT | N/A | 100% | 100 | N/A | N/A | 96% | 100 | N/A |
| 1 µM | N/A | 99% | 99 | 0.3149 | N/A | 100% | 100 | 0.0439 |
| 2 µM | N/A | 99% | 98 | 0.9943 | N/A | 96% | 100 | **0.0439** |
| 5 µM | N/A | 100% | 100 | 0.3124 | N/A | 100% | 100 | 0.0439 |
| 10 µM | N/A | 99% | 97 | 0.3099 | N/A | 99% | 100 | 0.3173 |
| 20 µM | N/A | 98% | 97 | 0.559 | N/A | 93% | 100 | **0.0306** |
| **Rose bengal** | | | | | | | | |
| **Concentration** | **Median Survival ‡** | **Endpoint Survival** | **n** | **Logrank *P* ¶** | **Median Survival ‡** | **Endpoint Survival** | **n** | **Logrank *P* ¶** |
| NT | N/A | 100% | 99 | N/A | N/A | 96% | 100 | N/A |
| 2 µM | N/A | 100% | 100 | N/A | N/A | 100% | 100 | 0.0559 |
| 5 µM | N/A | 100% | 99 | N/A | N/A | 99% | 100 | 0.3173 |
| 10 µM | N/A | 100% | 99 | 0.0987 | N/A | 97% | 100 | 0.3193 |
| 20 µM | N/A | 97% | 99 | 0.0987 | N/A | 99% | 100 | 0.2646 |
| 50 µM | N/A | 93% | 99 | 0.1897 | N/A | 100% | 100 | 0.3173 |
| 100 µM | N/A | 92% | 99 | 0.8045 | N/A | 95% | 100 | **0.0239** |

‡ N/A – not applicable because survival exceeded 50%

¶ P<0.05 are bolded

**Table S4. Comparison of larval survival between methylene blue and rose bengal following a photoperiod.** The survival of *Aedes aegypti* or *Anopheles gambiae* exposed to 2 μM, 5 μM, 10 μM, or 20 μM methylene blue and rose bengal were compared using the Log-Rank Mantel Cox test.

| ***Aedes aegypti*** | | | | | | | |
|---|---|---|---|---|---|---|---|
| | **Methylene blue** | | | ***Rose Bengal*** | | | |
| **Concentration** | **Median Survival** | **Endpoint Survival** | **n** | **Median Survival** ‡ | **Endpoint Survival** | **n** | **Logrank *P*** ¶ |
| 2 μM | 60 min | 7% | 97 | N/A | 92% | 100 | **<0.0001** |
| 5 μM | 40 min | 2% | 100 | N/A | 65% | 98 | **<0.0001** |
| 10 μM | 20 min | 0% | 99 | 24 hr | 44% | 95 | **<0.0001** |
| 20 μM | 20 min | 0% | 98 | 24 hr | 15% | 96 | **<0.0001** |
| ***Anopheles gambiae*** | | | | | | | |
| | **Methylene blue** | | | ***Rose Bengal*** | | | |
| **Concentration** | **Median Survival** | **Endpoint Survival** | **n** | **Median Survival** ‡ | **Endpoint Survival** | **n** | **Logrank *P*** ¶ |
| 2 μM | 24 hr | 16% | 100 | N/A | 99% | 99 | **<0.0001** |
| 5 μM | 60 min | 7% | 100 | N/A | 96% | 100 | **<0.0001** |
| 10 μM | 40 min | 2% | 100 | N/A | 87% | 100 | **<0.0001** |
| 20 μM | 30 min | 1% | 100 | N/A | 66% | 97 | **<0.0001** |

‡ N/A – not applicable because survival exceeded 50%
¶ P<0.05 are bolded

**Table S5. Comparison of *Aedes aegypti* and *Anopheles gambiae* survival following incubation with PSIs and a photoperiod.** The survival of *Ae. aegypti* and *An. gambiae* larvae following a 2 hr incubation with methylene blue and rose bengal and a photoperiod were compared using the Log-Rank Mantel Cox test.

**Methylene blue**

| | ***Aedes aegypti*** | | | ***Anopheles gambiae*** | | | |
|---|---|---|---|---|---|---|---|
| **Concentration** | **Median Survival ‡** | **Endpoint Survival** | **n** | **Median Survival ‡** | **Endpoint Survival** | **n** | **Logrank *P* ¶** |
| NT | N/A | 100% | 99 | N/A | 90% | 100 | 0.9811 |
| 1 µM | 24 hr | 34% | 98 | 24 hr | 33% | 100 | **0.0264** |
| 2 µM | 60 min | 7% | 97 | 24 hr | 16% | 100 | **<0.0001** |
| 5 µM | 40 min | 2% | 100 | 60 min | 7% | 100 | **0.0013** |
| 10 µM | 20 min | 0% | 99 | 40 min | 2% | 100 | 0.1478 |
| 20 µM | 20 min | 0% | 98 | 30 min | 1% | 100 | **0.0007** |

**Rose bengal**

| | ***Aedes aegypti*** | | | ***Anopheles gambiae*** | | | |
|---|---|---|---|---|---|---|---|
| **Concentration** | **Median Survival ‡** | **Endpoint Survival** | **n** | **Median Survival ‡** | **Endpoint Survival** | **n** | **Logrank *P* ¶** |
| NT | N/A | 90% | 99 | N/A | 90% | 100 | 0.9811 |
| 2 µM | N/A | 92% | 100 | N/A | 99% | 99 | **0.018** |
| 5 µM | N/A | 65% | 98 | N/A | 96% | 100 | **<0.0001** |
| 10 µM | 24 Hr | 44% | 95 | N/A | 87% | 100 | **<0.0001** |
| 20 µM | 24 Hr | 15% | 96 | N/A | 66% | 97 | **<0.0001** |
| 50 µM | 80 min | 1% | 98 | 120 min | 2% | 100 | **0.04** |
| 100 µM | 60 min | 0% | 100 | 80 min | 1% | 98 | 0.6331 |

‡ N/A – not applicable because survival exceeded 50%

¶ P<0.05 are bolded

**Table S6. Descriptive statistics of larvae following incubation in water and particulates.** Following a 2 hr incubation in water with (i) no particulates (NT), (ii) sand, or (iii) food, larvae were either transferred to a photoperiod (Light) or kept in darkness (Dark). Survival of larvae with either sand or food was compared between to the survival of larvae without any particulates (NT) using the Log-Rank Mantel Cox test; all significance indicates an increase in toxicity. NT, no treatment.

| | Light | | | | Dark | | | |
|---|---|---|---|---|---|---|---|---|
| ***Aedes aegypti*** | | | | | | | | |
| **Treatment** | **Median Survival ‡** | **Endpoint Survival** | **n** | **Logrank *P* ¶** | **Median Survival ‡** | **Endpoint Survival** | **n** | **Logrank *P* ¶** |
| NT | N/A | 99% | 80 | N/A | N/A | 99% | 80 | N/A |
| Food | N/A | 99% | 80 | >0.9999 | N/A | 100% | 80 | 0.3173 |
| Sand | N/A | 100% | 80 | 0.3173 | N/A | 100% | 80 | 0.3173 |
| ***Anopheles gambiae*** | | | | | | | | |
| **Treatment** | **Median Survival ‡** | **Endpoint Survival** | **n** | **Logrank *P* ¶** | **Median Survival ‡** | **Endpoint Survival** | **n** | **Logrank *P* ¶** |
| NT | N/A | 93% | 80 | N/A | N/A | 91% | 80 | N/A |
| Food | N/A | 99% | 80 | 0.054 | N/A | 98% | 80 | 0.0872 |
| Sand | N/A | 95% | 80 | 0.5309 | N/A | 98% | 80 | 0.0872 |

‡ N/A – not applicable because survival exceeded 50%

¶ P<0.05 are bolded

**Table S7. Descriptive statistics of larvae following incubation in methylene blue and particulates, without a photoperiod.** Following a 2 hr incubation in 2 μM or 20 μM methylene blue with either (i) no particulates (solution), (ii) sand, or (iii) food, larvae were maintained in darkness. Survival of larvae without any particulates (solution) was compared to the survival of larvae with sand or food using the Log-Rank Mantel Cox test.

| | ***Aedes aegypti*** | | | | ***Anopheles gambiae*** | | | |
|---|---|---|---|---|---|---|---|---|
| **2 μM Methylene blue** | | | | | | | | |
| **Treatment** | **Median Survival ‡** | **Endpoint Survival** | **n** | **Logrank *P* ¶** | **Median Survival ‡** | **Endpoint Survival** | **n** | **Logrank *P* ¶** |
| NT | N/A | 97% | 100 | N/A | N/A | 96% | 97 | N/A |
| Food | N/A | 99% | 100 | 0.3118 | N/A | 89% | 100 | 0.0741 |
| Sand | N/A | 98% | 100 | 0.6462 | N/A | 95% | 99 | 0.7779 |
| **20 μM Methylene blue** | | | | | | | | |
| **Treatment** | **Median Survival ‡** | **Endpoint Survival** | **n** | **Logrank *P* ¶** | **Median Survival ‡** | **Endpoint Survival** | **n** | **Logrank *P* ¶** |
| NT | N/A | 97% | 97 | N/A | N/A | 89% | 96 | N/A |
| Food | N/A | 97% | 99 | **0.0177** | N/A | 92% | 100 | 0.6658 |
| Sand | N/A | 98% | 99 | 0.6267 | N/A | 96% | 100 | 0.0502 |

‡ N/A – not applicable because survival exceeded 50%

¶ P<0.05 are bolded

**Table S8. Descriptive statistics of larvae following incubation in rose bengal and particulates, without a photoperiod.** Following a 2 hr incubation in 20 µM and 50 µM rose bengal with either (i) no particulates (solution), (ii) sand, or (iii) food, larvae were maintained in darkness. Survival of larvae without any particulates (solution) was compared to the survival of larvae with sand or food using the Log-Rank Mantel Cox test.

| | ***Aedes aegypti*** | | | | ***Anopheles gambiae*** | | | |
|---|---|---|---|---|---|---|---|---|
| **20 µM Rose bengal** | | | | | | | | |
| **Treatment** | **Median Survival ‡** | **Endpoint Survival** | **N** | **Logrank *P* ¶** | **Median Survival ‡** | **Endpoint Survival** | **n** | **Logrank *P* ¶** |
| NT | N/A | 98% | 98 | N/A | N/A | 97% | 98 | N/A |
| Food | N/A | 98% | 97 | 0.9918 | N/A | 95% | 100 | 0.7587 |
| Sand | N/A | 100% | 98 | 0.1562 | N/A | 96% | 100 | 0.9768 |
| **50 µM Rose bengal** | | | | | | | | |
| **Treatment** | **Median Survival ‡** | **Endpoint Survival** | **n** | **Logrank *P* ¶** | **Median Survival ‡** | **Endpoint Survival** | **n** | **Logrank *P* ¶** |
| NT | N/A | 8% | 90 | N/A | N/A | 96% | 96 | N/A |
| Food | N/A | 100% | 90 | **0.0071** | N/A | 95% | 100 | 0.4848 |
| Sand | N/A | 97% | 90 | 0.1943 | N/A | 97% | 98 | 0.1759 |

‡ N/A – not applicable because survival exceeded 50%
¶ P<0.05 are bolded

**Table S9. Descriptive statistics of larvae following incubation in methylene blue and particulates, with a photoperiod.** Following a 2 hr incubation in 2 μM or 20 μM methylene blue with either (i) no particulates (solution), (ii) sand, or (iii) food, larvae were transferred to a photoperiod. Survival of larvae without any particulates (solution) was compared to the survival of larvae with sand or food using the Log-Rank Mantel Cox test.

| | *Aedes aegypti* | | | | *Anopheles gambiae* | | | |
|---|---|---|---|---|---|---|---|---|
| **2 μM Methylene blue** | | | | | | | | |
| **Treatment** | **Median Survival** | **Endpoint Survival** | **n** | **Logrank *P* ¶** | **Median Survival ‡** | **Endpoint Survival** | **n** | **Logrank *P* ¶** |
| NT | 60 min | 4% | 100 | N/A | 24 hr | 19% | 98 | N/A |
| Food | 110 min | 21% | 100 | **<0.0001** | N/A | 59% | 99 | **<0.0001** |
| Sand | 60 min | 1% | 98 | 0.3051 | 24 hr | 29% | 100 | **0.0295** |
| **20 μM Methylene blue** | | | | | | | | |
| **Treatment** | **Median Survival** | **Endpoint Survival** | **n** | **Logrank *P* ¶** | **Median Survival** | **Endpoint Survival** | **n** | **Logrank *P* ¶** |
| NT | 40 min | 1% | 97 | N/A | 120 min | 15% | 98 | N/A |
| Food | 20 min | 0% | 98 | **0.0207** | 40 min | 1% | 99 | **<0.0001** |
| Sand | 20 min | 1% | 96 | 0.0662 | 120 min | 12% | 98 | 0.5738 |

‡ N/A – not applicable because survival exceeded 50%

¶ P<0.05 are bolded

**Table S10. Descriptive statistics of larvae following incubation in methylene blue and particulates, with a photoperiod.** Following a 2 hr incubation in 20 µM or 50 µM rose bengal with either (i) no particulates (solution), (ii) sand, or (iii) food, larvae were transferred to a photoperiod. Survival of larvae without any particulates (solution) was compared to the survival of larvae with sand or food using the Log-Rank Mantel Cox test.

| | *Aedes aegypti* | | | | *Anopheles gambiae* | | | |
|---|---|---|---|---|---|---|---|---|
| **20 µM Rose bengal** | | | | | | | | |
| **Treatment** | **Median Survival** | **Endpoint Survival** | **n** | **Logrank *P*** ¶ | **Median Survival** | **Endpoint Survival** | **n** | **Logrank *P*** ¶ |
| NT | 24 hr | 35% | 97 | N/A | 24 hr | 32% | 95 | N/A |
| Food | 40 min | 0% | 95 | **<0.0001** | 24 hr | 39% | 99 | 0.6149 |
| Sand | 120 min | 8% | 97 | **<0.0001** | 24 hr | 22% | 99 | 0.6503 |
| **50 µM Rose bengal** | | | | | | | | |
| **Treatment** | **Median Survival** | **Endpoint Survival** | **n** | **Logrank *P*** ¶ | **Median Survival** | **Endpoint Survival** | **n** | **Logrank *P*** ¶ |
| NT | 80 min | 3% | 97 | N/A | 100 min | 2% | 98 | N/A |
| Food | 40 min | 0% | 98 | **<0.0001** | 80 min | 5% | 100 | **0.0384** |
| Sand | 40 min | 1% | 100 | **<0.0001** | 120 min | 8% | 97 | 0.1528 |

¶ P<0.05 are bolded

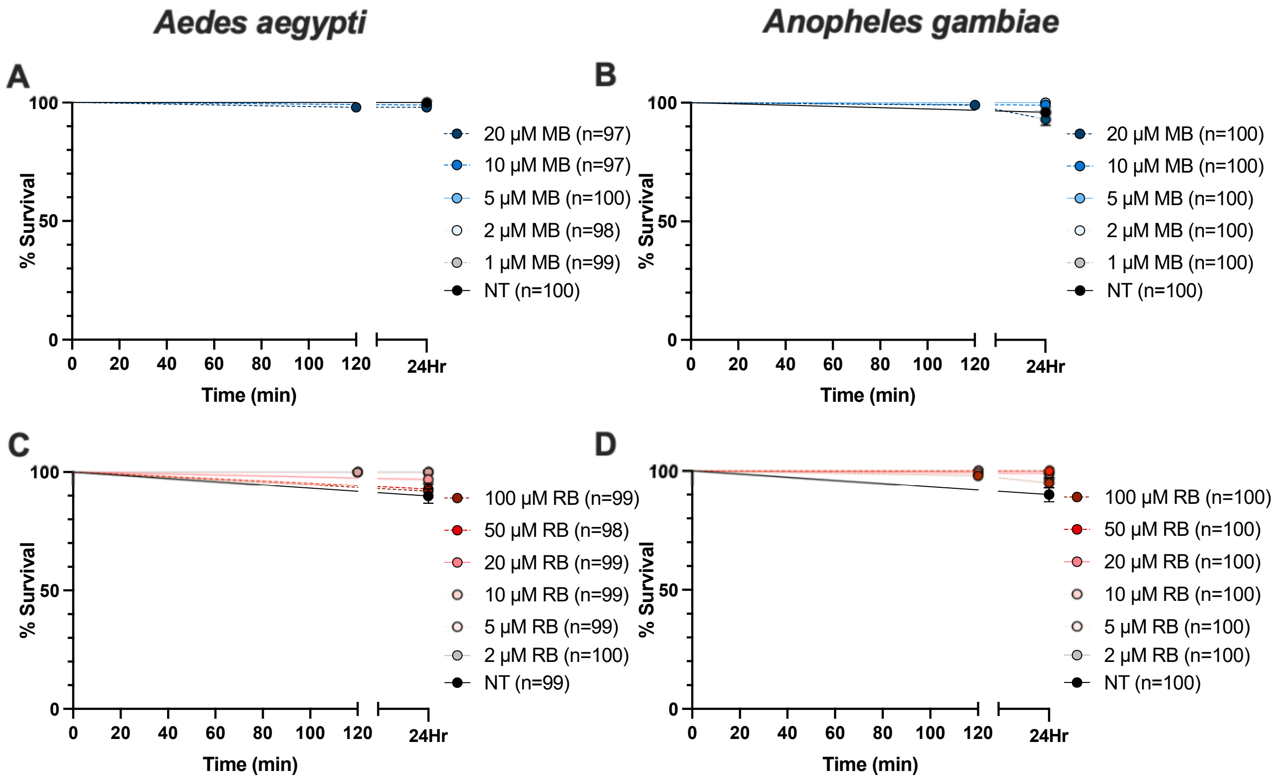


**Figure S1.** Larval survival following incubation with various concentrations of PSIs in the dark. **(A-D)** Survival of *Ae. aegypti* (A, C) and *An. gambiae* (B, D) following 2 hr incubation in various concentrations of methylene blue (A, B) and rose bengal (C, D). Whiskers indicate the standard deviation MB, methylene blue; RB, rose bengal; NT, no treatment.

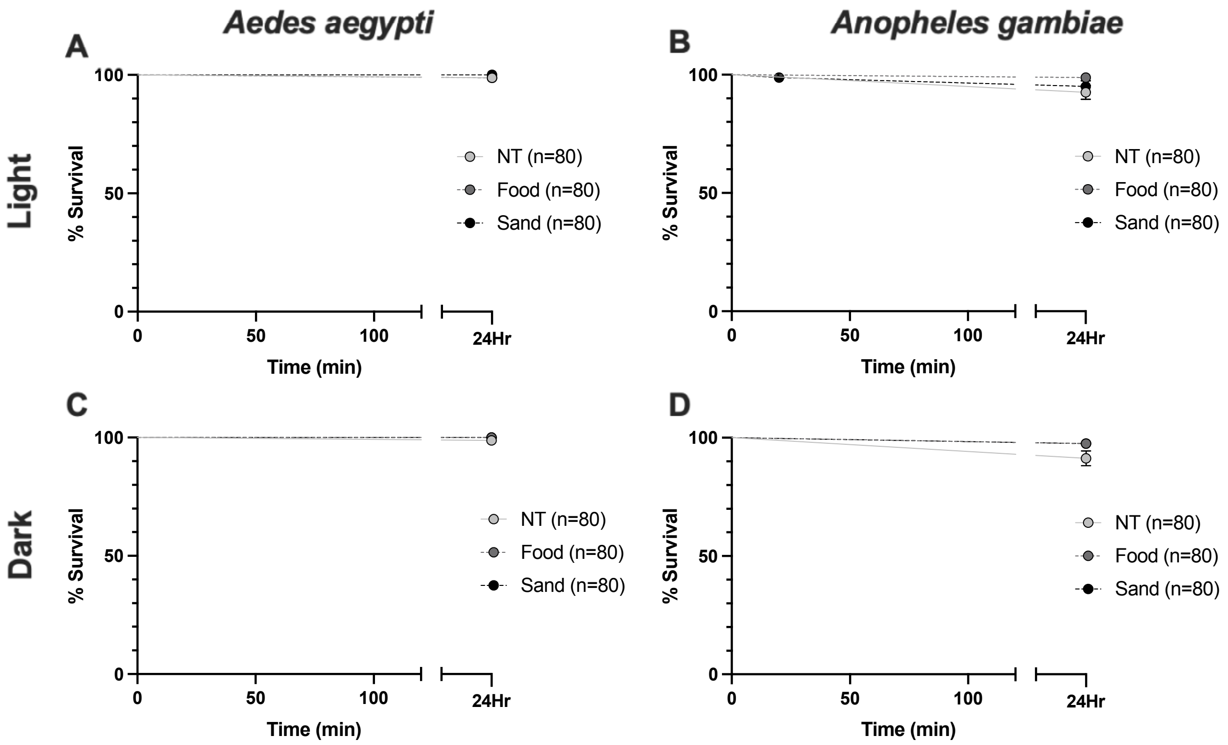


**Figure S2.** Larval survival in the presence of sand or food in the dark. Survival of **A.** *Ae. aegypti* and **B.** *An. gambiae* maintained in darkness following a 2 hr incubation in water without particulates (solution), with food, or with sand. Survival of **C.** *Ae. aegypti* and **D.** *An. gambiae* maintained in darkness following a 2 hr incubation water without particulates (solution), with food, or with sand. Whiskers indicate the standard deviation. NT, No treatment.

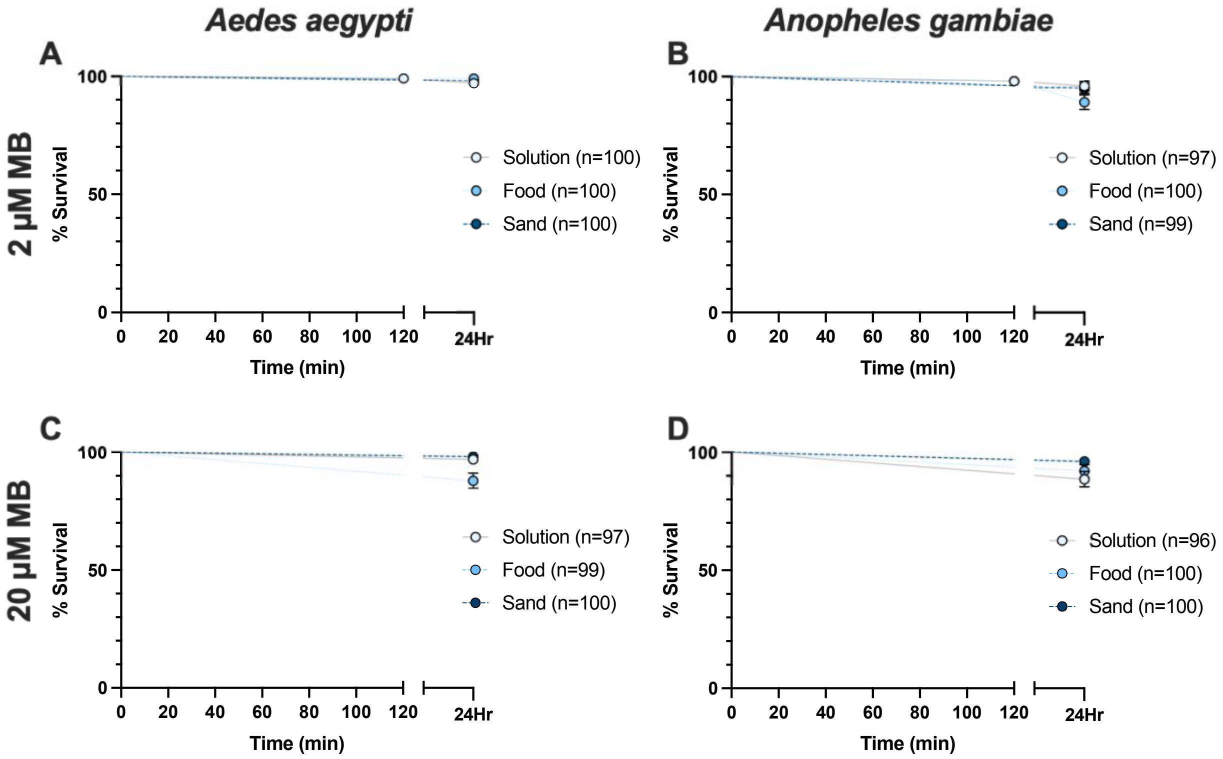


**Figure S3.** Survival of larvae incubated in methylene blue and particulates, without a photoperiod. **(A-D)** Survival of *Ae. aegypti* (A, C) and *An. gambiae* (B, D) throughout a 2 hr photoperiod following a 2 hr incubation in 2 µM methylene blue (A, B) or 20 µM methylene blue (C, D) without particulates (solution), with food, or with sand. Whiskers indicate the standard deviation. MB, methylene blue.

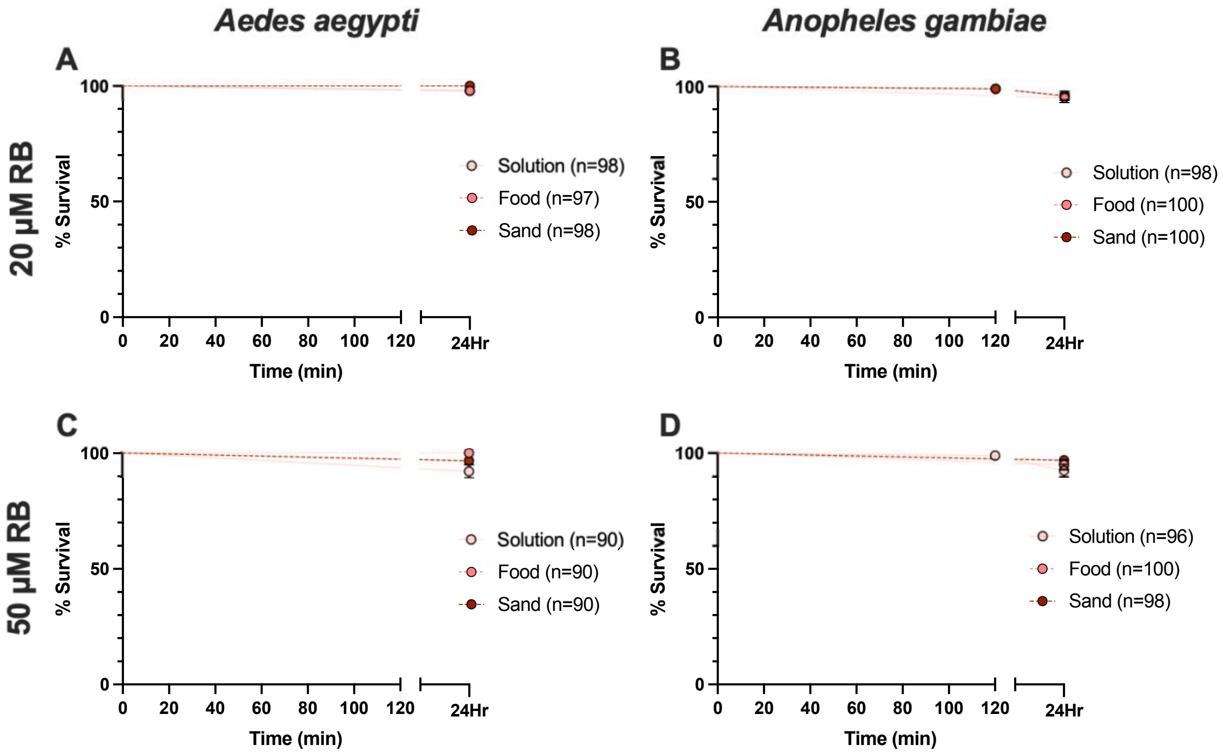


**Figure S4.** Survival of larvae incubated in rose bengal and particulates, without a photoperiod. **(A-D)** Survival of *Ae. aegypti* (A, C) and *An. gambiae* (B, D) throughout a 2 hr photoperiod following a 2 hr incubation in 20 μM rose bengal (A, B) or 50 μM rose bengal (C, D) without particulates (solution), with food, or with sand. Whiskers indicate the standard deviation. RB, rose bengal.
